# Randomly oriented microgrooved hydrogel guides cellular motility, modulates speed, and governs directionality of cellular spread

**DOI:** 10.1101/2024.09.10.612339

**Authors:** Biswajoy Ghosh, Krishna Agarwal

**Affiliations:** Department of Physics and Technology, UiT-The Arctic University of Norway, Tromsø, Norway

## Abstract

Cell migration is a fundamental biological process, yet the mechanisms underlying how cells sense and navigate complex environments remain poorly understood. In this study, we developed a system of randomly oriented microgrooves, designed at cellular length scales, to explore motility intelligence in response to varied topographies. These microgrooves allowed cells to freely choose their migratory paths, revealing key insights into how cells sense and adapt to topological cues. Using fibroblast cells migrating over these grooved substrates, we examined cellular processes such as actin cytoskeleton remodeling, cell adhesion dynamics, and the impact of groove alignment on migration speed and directionality. Our results demonstrate that cells align their cytoskeletal structures to groove geometries, forming actin-rich anchors that enhance migration in groove-aligned environments. Cells migrating in grooves aligned with their intrinsic polarity exhibited faster, more directed migration compared to those in misaligned or control conditions. This work advances our understanding of cell-topology interaction and provides new perspectives for tissue engineering applications in cancer therapy and wound healing.

## INTRODUCTION

Cell migration is a fundamental process in biological systems, serving as the basis for wound healing, immune responses, cancer metastasis, and evaluating drug efficacy. This dynamic behavior involves changes cell morphology at multiple scales in response to environmental cues. This makes understanding cell migration difficult limiting possibilities to harness the traits for applications in healthcare.

Cellular intelligence, which governs migration, is closely linked to the cells’ sensing and reaction mechanisms. These mechanisms are fundamental to various types of motility, including chemotaxis, topotaxis, geotaxis, haptotaxis, and durotaxis. A key aspect of cellular intelligence is the ability to make decisions and move toward specific goals. Among the different types of movement, cells frequently navigate textured surfaces and obstacles within the body, a process known as topotaxis. Park et al.^1^ reintroduced the concept of topotaxis, demonstrating that cells migrate in response to nanoscale topographic gradients within their extracellular matrix (ECM). Their work showed that substrate topology significantly influences cell motility.

Historical studies on cell migration have laid the foundation for understanding how cells interact with their surroundings. Early research emphasized the role of protein adhesion to substrates^2^ and the effects of shear stress on ion channels^3^. These studies provided early insights into how cells respond to mechanical forces. Further discoveries, such as the formation of fibroblast microfilament bundles synchronizing with focal adhesions^4^, highlighted the connection between cellular structures and their migration. The discovery of integrins, a protein family critical for cellular attachment to substrates and sensory mechanisms, advanced the understanding of mechanotransduction, wherein cells sense and react to their mechanical environment^4^.

Topotaxis, which involves cell guidance by topographical features such as collagen fibers and ECM pores^5–7^, plays a key role in tissue organization and disease progression. Leukocytes migrate through ECM trails during immune responses without ECM remodeling^8,9^, while cancer cells often remodel ECM proteolytically or deform it to create migration paths^10^. Multiscale hierarchical topographies inspired by collagen architecture have been shown to influence stem cell morphology, orientation, and differentiation, particularly enhancing osteogenesis^11–13^. Cancer cells can orient themselves parallel to the surrounding tissue’s topological features and migrate directionally without significant ECM remodeling, indicating their ability to sense complex microenvironmental structures and adjust their migration accordingly^14^.

Substrate texture has been shown to affect cell function that is cell differentiation and maturation using a range of materials and techniques. For example, tissue mimicking hydrogels with tunable mechanical properties^15–31^ have been studied for different cell types to understand the proliferation, differentiation and other cellular functions. Stem cell differentiation is optimized by controlling hydrogel stiffness, with specific ranges promoting cardiomyocytes, osteoblasts, or adipocytes^32^. Decellularized ECMs provide a 3D structure supporting tissue-specific cell differentiation^33^. Thin-film hydrogels are commonly used, such as graphene-coated PDMS of varying stiffness, which enhanced MSC osteogenesis and expression of osteogenic markers and integrin/FAK proteins compared to PDMS alone, regardless of stiffness^34^. Cellular traction forces on thin silicon film substrates were studied by analyzing elastic distortion and wrinkling^35^. A natural polyisoprene membrane promoted ADMSC differentiation into neural precursors without induction factors, likely through mechanotransduction involving YAP and AMOT proteins^36^. Surface patterns and architectural cues also influence cell behavior and differentiation. Microprinting^37,38^ and micropatterning^39–43^ create surface patterns on substrates like PEG hydrogels and polyimide polymers, which enhance MSC differentiation. PDMS substrates with microgrooves promote cell elongation and tenogenic marker expression^44^, while microgrooves and scaffold geometry on PLGA thin films enhance differentiation towards myocardial lineages and cardiomyocytes^45,46^. V-shaped ridges on fibrin gels align cardiac tissue^47^, and scaffold geometry affects cardiovascular tissue growth and remodeling^48^. Nanostructures, like nanopillars and nanotubes, induce osteogenesis, while nanofiber orientations in scaffolds improve cell behavior in nerve, ligament, and tendon regeneration^49–55^. Nanopits fabricated on various polymers aid in studying cell behavior^56–59^. Nanoscale grooves and pillars influence stem cell morphology and differentiation through mechanotransduction^60–63^. Nanofiber orientations further enhance regeneration in nerve, ligament, tendon, and bone tissues, with aligned fibers supporting tendon and cardiac regeneration^64–70^. Scaffold stiffness, 3D geometry, and fiber diameter also influence differentiation, with larger fibers enhancing tenogenesis and 3D electrospun scaffolds promoting adipogenesis.

Despite these advances, the key aspect of cell migration remains elusive: the understanding of cellular intelligence. Traditional research has focused on using materials as substrates to observe cell movement, but not to uncover the underlying decision-making processes. To address this, we created randomly oriented microgrooves, designed at cellular scales (tens of microns), which are arranged in multiple directions. These microgrooves allow cells to navigate freely and choose their migration paths, offering insights into their decision-making and directional preferences. In this study, we aim to explore the basis of migratory intelligence by investigating the underlying cellular processes—particularly changes in actin, the cytoskeleton, and cell adhesion—when cells navigate these microgroove mazes, both collectively and individually. Through this approach, we seek to better understand how cells sense and respond to complex environments.

## RESULTS

### Groove patterned soft hydrogels promote collective cell migration activity

We prepared grooved hydrogels with the stiffness of 1.4 ± 0.5*kPa*. The hydrogel had different shapes like straight, zig-zag, y-forked, etc., and was present as a 200 − 300*µm* patch. The cells were seeded right before the patch and they had to migrate after crossing the grooved patch to reach the plane soft hydrogel (Figure 1a). The grooves had a mean thickness of 2.1*µm* (Figure 1b) and presented angles uniformly distributed from 0 to 90^°^*C* relative to the line of the patch (Figure 1c). We allowed the fibroblast cells to migrate across the patch (Figure 1d). The cells successfully traversed the patch for a total of 60 hours (Figure 1h). To investigate the significant time taken by the cells to cross this patch, we observed the standard deviation maps of cell migration in three 6-hour windows in the beginning, middle, and near the end of the migration task, represented by Figure 1 e, f, and g, respectively. We observed a high amount of standard deviation concentrated in the groove patch, indicating high cellular activity in the patch region in comparison to the region before the patch. Further, even when the cells were close to the end of the texture patch (Figure 1g), the standard deviation at the beginning of the texture continued to have high activity. Upon closer investigation, we observed that the cells were aligned and migrated along the direction of the grooves (Figure 1i). We generated temporal color maps of cellular activity in the middle and the end of the cells’ collective migration across the patch (Figure 1j and k). When the cells were in the middle of the patch (Figure 1j), the leading edge of the cells showed prominent blobs (yellow arrows) indicating high activity in navigating the grooves and exploring multiple directions. At the end of the collective migration patch (Figure 1k), these high activity blobs were not observed any longer. Further, as the cells completed traveling across the patch, a shift in the line that marked the end of the texture was observed. This implies traction force generated by the collective cell migration causes the pull effect of the texture.

**Figure 1.**
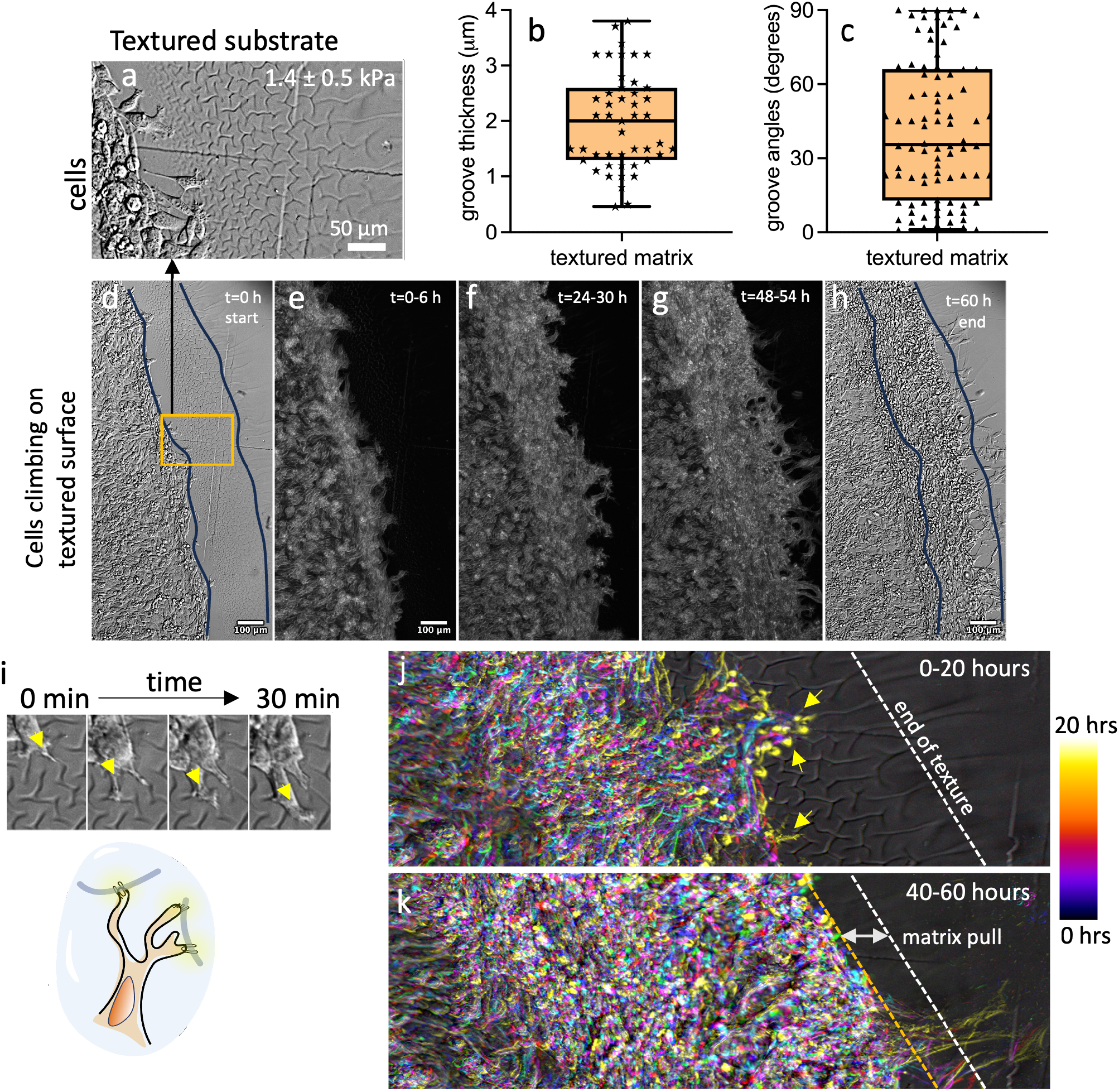
Groove-textured hydrogel promotes directed collective cell migration. (a) shows a view of fibroblast cells starting to climb the groove-textured patch made of soft GelMA hydrogel under a Differential interference contrast (DIC) microscope. (b) distribution of the thickness of the grooves, (c) angular distribution of the grooves relative to the line of texture. (d) cells initiate motility towards the texture. (e-g) standard deviation maps showing most active motility regions on the grooves textured patch in different 6-hour windows. (h) cells complete traveling through the groove-textured patch. (i) shows thin cellular processes sensing grooves and moving forward in a 30-minute window. The diagrammatic illustration below represents the cellular processes to anchor into grooves to move forward. (j) color map showing the high activity of leading cells when the cells are moving in the grooves in a 20-hour window. The white dashed line marks the end of the grooved patch. (k) color map showing reduced activity of leading cells after cells have traversed the grooved patch. The yellow dashed line marks the shift in the end of the grooved patch.

### Cell cytoskeleton acquires groove morphology to move

We investigated how the cells’ cytoskeleton in their leading edges accommodated the grooves in the substrate (Figure 2a). When the grooves have a Y-shape, the actin in the leading edges acquires the bifurcating morphology to explore both the grooved directions (Figure 2b and c). We performed Super-Resolution Radial Fluctuations (SRRF) microscopy (abbreviated as SR) to resolve actin nanostructures that are otherwise not visible with a confocal fluorescence microscope (can resolve up to 50 nm). However, the cells eventually choose one of the two parts and allocate more actin polymerization and bundling to one of the two paths (Figure 2d and e). This is further illustrated schematically in (Figure 2f). The leading edge of a cell moving on the surface of a smooth hydrogel (Figure 2g) does not have a single narrow and thick bundle of actin on its leading edge (Figure 2h). Instead, it consists of hundreds of actin filaments distributed on the leading edge (Figure 2i and j). We further observe cell motility along a straight groove (Figure 2k and m). The actin was bundled along a straight groove into thick bundles of 100-150 nm (Figure 2 n, o, and p). Figure 2i illustrates this aligning of the actin bundles along the grooves. Besides sharp bifurcation and straight lines, we observed cell motility in curved grooves (Figure 2q). Migrating cells adjust their shape to the geometry of the surrounding substrate, to migrate in a preferred direction^71–73^. We found the actin almost completely aligned itself to the curvature of the groove (Figure 2r). The cell following also exhibited actin shaping based on the curvature sensing (as shown by the yellow arrow in Figure 2r). Although the actin thick bundle formed across the length of the cell, a small bifurcation of the actin was observed at the leading edge (Figure 2s) as the cell reached the vertex of a bifurcating groove. The two thick bundles at the leading edge were roughly 10*µm* in length into the cell, implying rapid actin bundling and polymerization based on topology sensing upon encountering the bifurcation.

**Figure 2.**
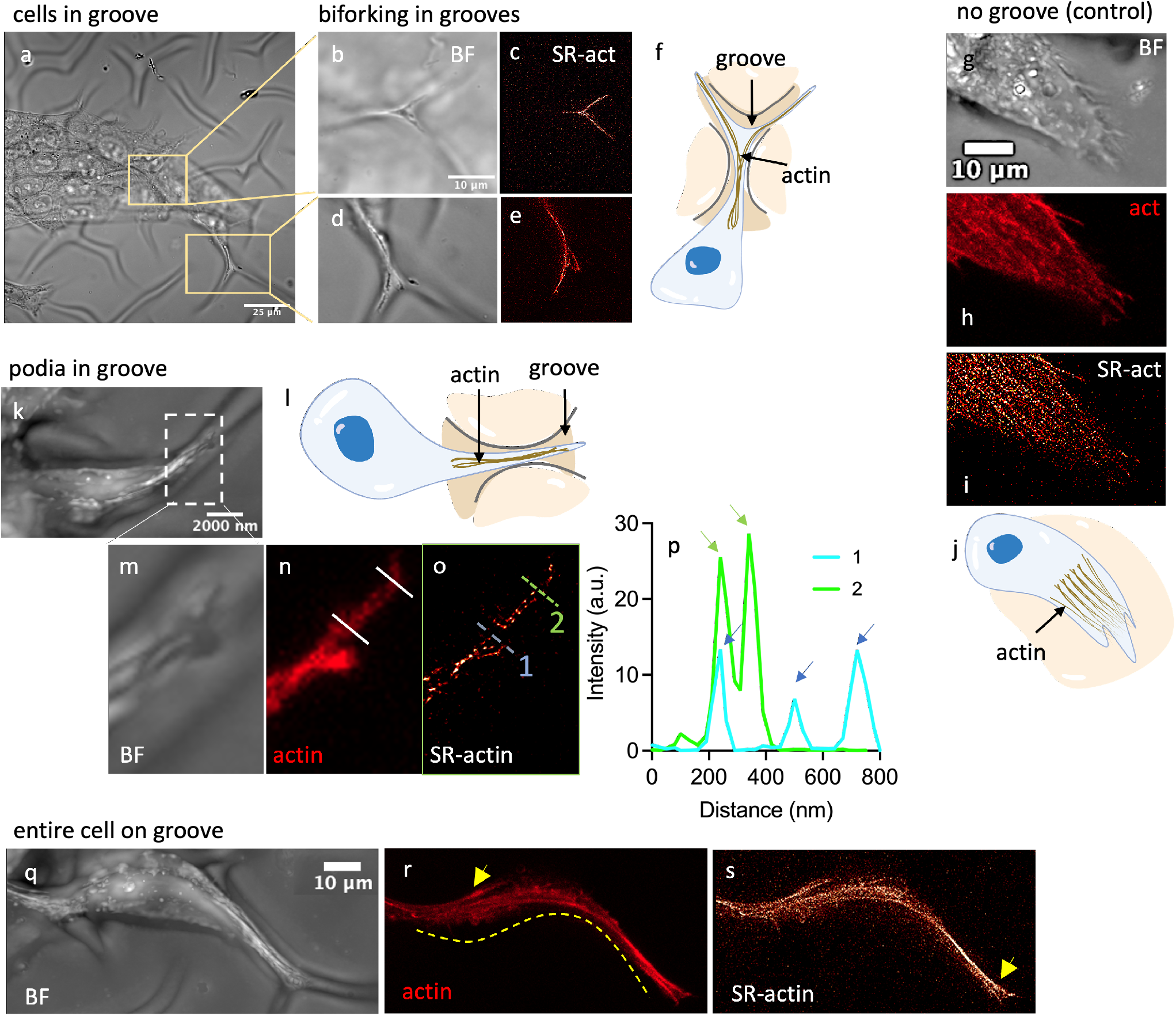
Actin organization and dynamism during cell migration on hydrogel grooves. (a) bright-field (BF) image of the cells moving on the grooved texture. (b,d) insets showing two regions where cells migrate at a Y-shaped groove, bifurcating in two directions and the leading edge of the cell following both grooves. (c,e) Super-Resolution Radial Fluctuations (SRRF) microscopy (SR) images of actin show thick bundling of actin following the bifurcation of the groove. (f) shows a diagrammatic image of actin response to a Y-shaped groove. (g,h) BF and actin images (confocal microscopy) of the leading edge of the cell moving on a flat hydrogel (control sample). (i) SR image shows many thin actin fibers. (j) simple illustration of showing actin organization on the leading edge of the cell moving on a non-grooved control hydrogel. (k) the leading edge of a cell moving into a straight groove aligned with the directional polarity of the cell. (l) illustration showing how the cell squeezes into the grooves. (m,n) insets of BF and actin images of the podial structure inside the groove. (o) SR images showing multiple actin bundles. (p) plot shows two profiles along the podia (see dashed lines 1 and 2 in (o) of which the profile was created. The dashed lines correspond spatially to the solid white lines in (n) to indicate position) with peaks of ∼100 nm width and each peak resembling a bundle. (q) BF image of a cell moving on a curved groove. (r) actin image showing the maximum projection of the lower half of the cell in contact with the groove acquires the shape of the groove. The yellow arrow shows the leading edge of a cell behind following the same track. (s) SR image of the lower half of the cell shows thick bundling of the actin fibers across the entire length of the cell. The yellow arrow shows the forking of the actin bundles as it is about to approach a Y-shaped groove.

### Actin polymerization and cell-matrix adhesion co-occur in cellular podia in the grooves

Since motility is governed by the actin polymerization and focal adhesion complexes that attach the cells to the matrix via integrins, we investigated spatial localization of actin cytoskeleton and cell-matrix adhesion protein vinculin (Figure 3a-c). We found that the density of the cytoskeleton was proportional to the density of distribution of vinculin (Figure 3b and c insets). As the actin and vinculin together function in leading edge protrusion and traction force generation, the two molecules are present in the cells attached to the grooves (Figure 3d). We also observed the co-localization of the two proteins if the shape of the groove was curvy (Figure 3e, f, and g). We found a strong co-localization between the two proteins (Figure 3h).

**Figure 3.**
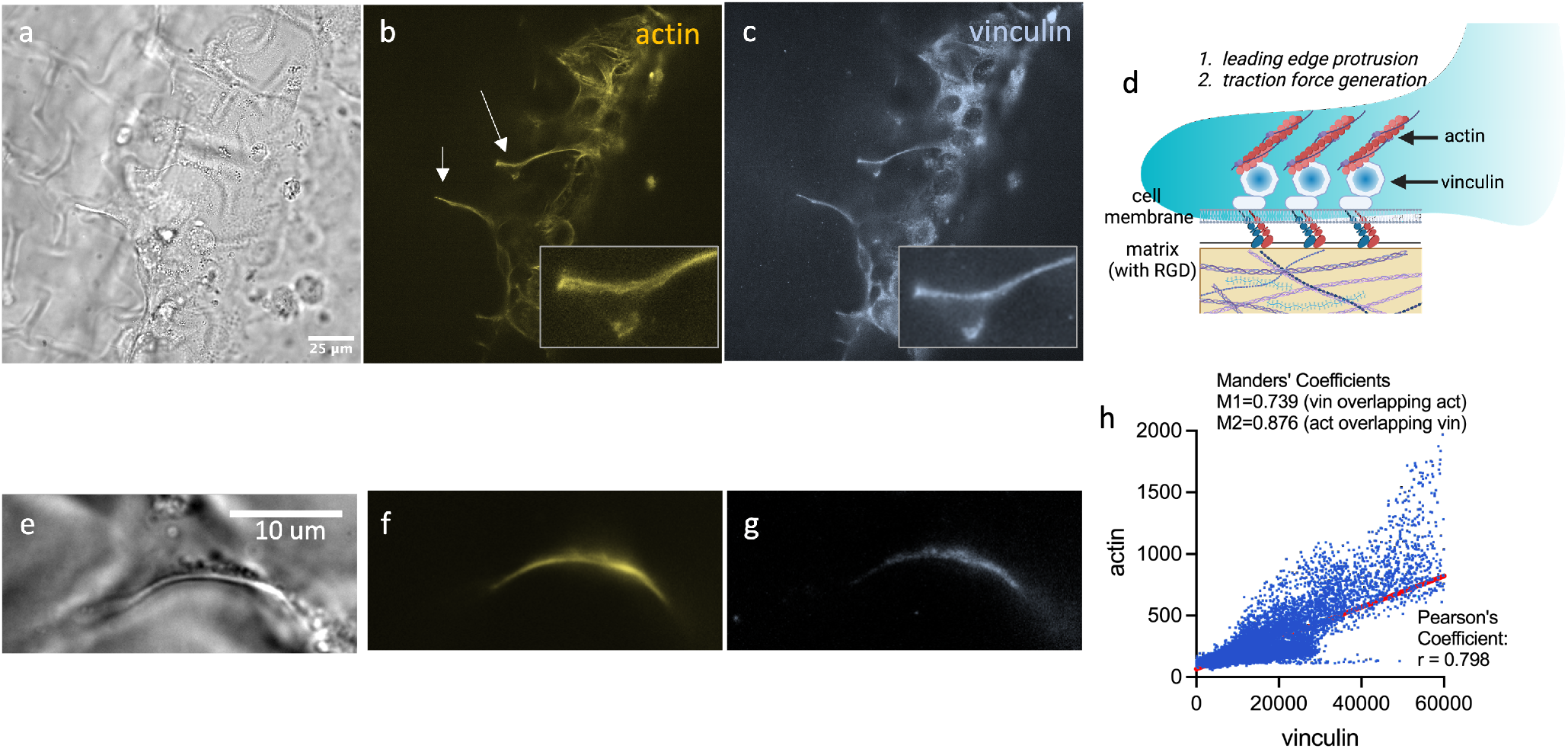
Co-localization of the cytoskeleton and cell adhesion markers of cellular processes in grooves. (a) BF image of a full field showing cells on grooves. (b,c) actin and focal adhesion protein vinculin. Arrows show long cellular podial processes in the grooves. Insets in (b and c) show the close-up of the spatial distribution of the proteins in a y-shaped groove with one of the bifurcating arms having a higher density of proteins. (d) illustration showing how actin and vinculin function together for cell-matrix adhesion. (e-g) BF, actin, and vinculin images of a cell on the curved groove. (h) a cytofluorogram showing the co-localization of actin and vinculin in (f and g) with Pearson’s correlation coefficient of 0.798 indicating significant co-localization.

### The actin reorganization occurs in different planes of the cell indicating dynamics of cell-groove interaction

Since actin organization is a dynamic process, we investigated live changes in unlabelled cells as they navigated the grooves. Gradient Light Interference Microscopy (GLIM)^74–76^ is a powerful tool to observe high-resolution structural changes in different depth planes of the cell (Figure 4). The intensity in the GLIM images represents the gradient phase which is a measure of cells solid mass. We used two distinct planes of the cells to visualize the cell-groove interaction - the groove plane and the surface plane. The analysis reveals the distinct behavior of the cytoskeleton in both the surface plane and the groove plane over time. At time = 0, the in-groove part of the cell (Figure 4e) shows a clear thick cytoskeletal bundle (bounded by dashed lines and white arrow). A part of the thickened cytoskeleton bundle is also visible in the surface plane (Figure 4a). At time = 4 minutes, the groove plane (Figure 4f) displays a bifurcation of the cytoskeleton, with two distinct arms indicated by white and yellow arrows. In contrast, no significant changes are observed in the surface plane (Figure 4b) at this time point. By time = 8 minutes, the groove plane (Figure 4g) shows a clear preference of the leading edge towards one direction (white arrow), and a small protrusion is noted on the rear side (blue arrow), suggesting podial exploration. Concurrently, the surface plane (Figure 4c) reveals an increase in cytoskeletal bundling on the side edge (blue arrow), correlating with the observed podial exploration. At time = 12 minutes, the groove plane (Figure 4h) demonstrates further movement of the cell towards the preferred direction (white arrow). However, a small bump remains on the other arm of the fork (yellow arrow), and on the side rear (blue arrow), indicating potential retraction in that direction. On the surface plane (Figure 4d), two distinct blunt knots are observed—one on the side rear (blue arrow) and another along the direction of the alternate fork path (yellow arrow). The gradient phase value (Figure 4i) indicates the cells’ dry mass concentration, showing a significantly higher value in the groove plane. This suggests a higher density of cytoskeletal structures within the groove compared to the surface plane. The changes in the grooves correspond to cytoskeleton changes in the surface plane, implying a 3D reorganization of actin that spans the height of the cell.

**Figure 4.**
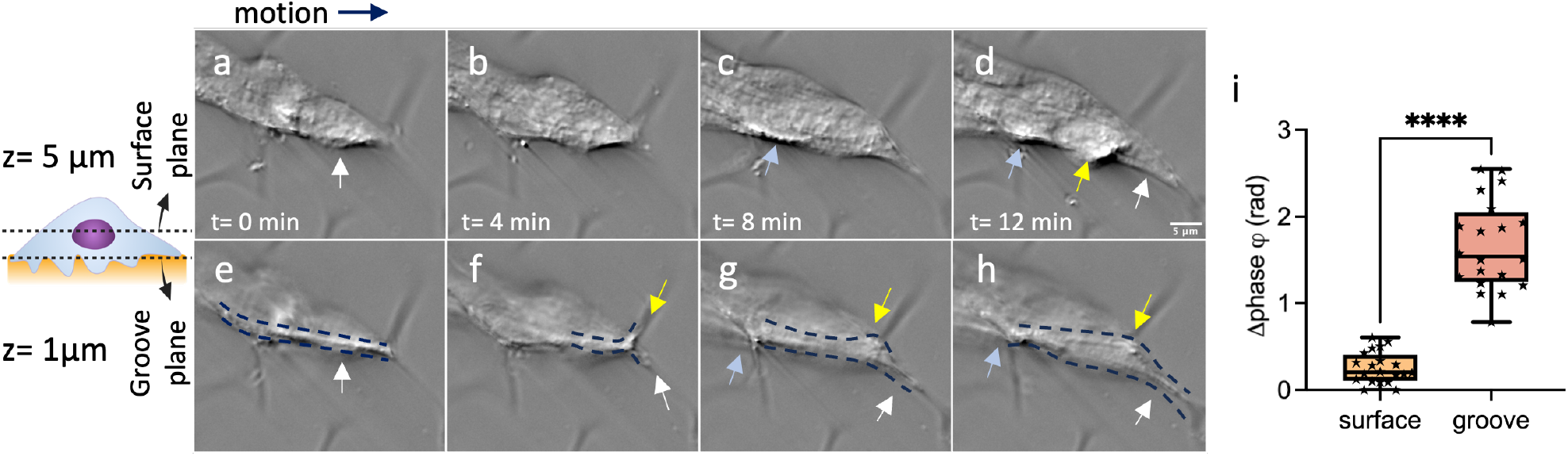
Label-free cytoskeletal dynamics at different depth planes of the cells on groove using Gradient Light Interference Microscopy (GLIM). (a-d) images of cells moving on grooves in the top half of the cell **(surface plane)** which lies above the groove. (e-h) the corresponding bottom half of the cell **(groove plane)** including the parts embedded in the groove. (a,e) at time = 0, the in-groove part of the cell shows a clear thick cytoskeletal bundle (bounded by dashed lines and white arrow). A part of the thickened cytoskeleton bundle is also visible in the surface plane. (b,f) at time = 4 mins, the groove plane witnesses a bifurcation indicated by the white and yellow arrows representing the two arms. In the surface plane, no prominent changes are visible. (c,g) at time = 8 mins, in the groove plane, the leading edge makes a clear preference towards one direction (white arrow). Further, a small protrusion is observed on the rear side in the grooved plane (blue arrow) indicating podial exploration. on the surface plane, the side edge witnessed an increase in the cytoskeletal bundling (blue arrow) corresponding to the podial exploration. (d, h) at time = 12 mins, in the groove plane, the cell further moves towards the preferred direction (white arrow). However, a small bump is still observed on the other arm of the fork (yellow arrow), and on the side rear (blue arrow) indicating possible retraction of the cell leading in that direction. On the surface plane, two blunt knots are visible on the side rear (blue arrow) and along the direction of the alternate fork path (yellow arrow). (i) shows the gradient phase value indicative of the cells’ dry mass concentration. It shows a significantly higher value in the grooved plane indicating a high density of cytoskeletal structures.

### Alignment of the groove relative to the cell’s polarity affects speed of cell motility

Figure 5 illustrates the effect of groove orientation relative to cell migration polarity on cell speed and directionality. When grooves align with cell polarity (Figure 5a), cells move faster, as indicated by the motion profile of the cell’s centroid (blue dashed line). Conversely, when grooves are nearly perpendicular to the cell’s migration polarity (Figure 5b), multiple podial structures form at the leading edge, slowing down the cell’s speed. The motion profile (blue dashed line) confirms reduced migration efficiency. Figure 5c quantifies cell speed in control samples (plane soft hydrogels), grooves aligned with cell polarity, and grooves against cell polarity. The schematic in Figure 5d explains that aligned grooves result in condensed, directed actin bundles, generating higher traction forces and thus increasing speed, while multi-directional podia in misaligned grooves produce lower traction forces, reducing speed. Our observations align with previous cell migration studies reported in the literature^77^. We observed that the speed of the cells moving in the grooves aligned with cell polarity was almost twice the speed of the cells in the control sample (Figure 5c). This led us to investigate the possibility of additional mechanisms that can increase cellular speed in grooves aligned with the cell polarity.

**Figure 5.**
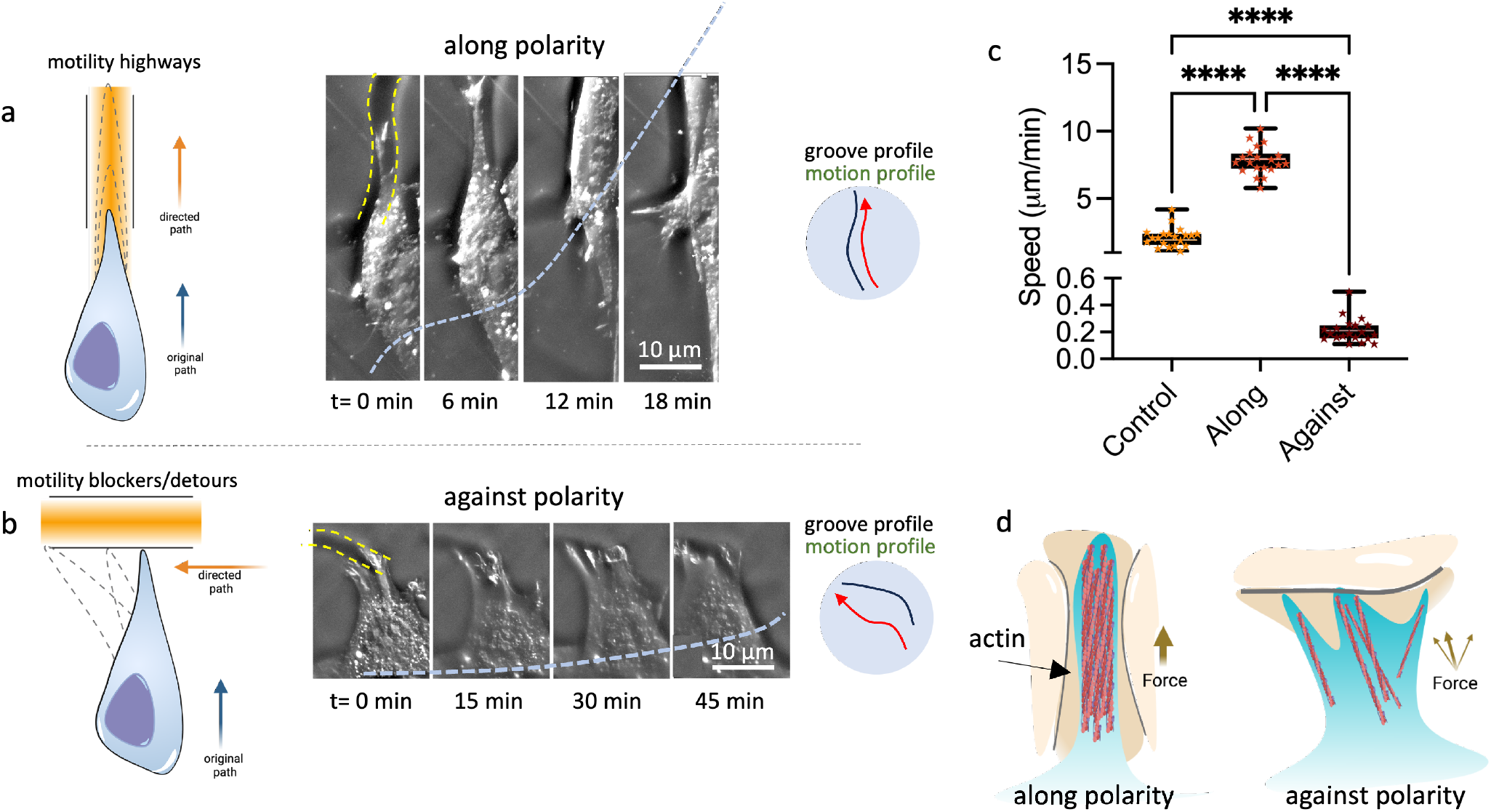
Influence of groove direction relative to cell migration polarity on cellular migration speed and directionality. (a) shows a case where the grooves are aligned with the cell polarity which makes cells move faster. The blue dashed line indicates the motion profile of the cell’s centroid. (b) another case where the groove is aligned nearly perpendicular to the direction of the cell’s migration polarity causing multiple podial structures to erupt from the leading edge of the cell, reducing the speed. The dashed blue line indicates the motion profile of the cell’s centroid. (c) speed of cells in control (plane soft hydrogels), groove along the cell polarity, and the groove against cell polarity. (d) An illustration to show a higher force generation in condensed and directed actin bundle in an aligned groove, compared to lower traction force generated by multi-directional podia in the grooves against the direction of polarity.

### Cells develop anchoring spikes of actin in the grooves to propel cells forward

To unravel additional mechanisms of regulating speed in grooves, we observed 3D rendered actin organization within the grooves. On flat, soft hydrogel surfaces (control conditions), cells exhibited blunt leading edges with no prominent actin bundling (Figure 6a). A 3D-rendered side view of these cells revealed a thin layer of cytoplasm with limited active bundling, indicating minimal engagement with the substrate (Figure 6b). In contrast, cells migrating on grooved hydrogel surfaces displayed concentrated leading edges that actively aligned within the grooves (Figure 6c). The 3D side view perspective demonstrated that these cells formed pronounced anchor-like actin processes, which were deeply embedded in the grooves (Figure 6d, yellow arrows). The bottom view further highlighted these actin anchors, which were oriented along the groove structures in various directions (Figure 6e). A cross-sectional view of the leading edge revealed multiple actin-based anchors, indicating a robust interaction between the cell and the grooved substrate (Figure 6f). A histogram of the angles of these actin anchors with respect to the direction of cell migration indicated a broad distribution. However, there was a clear dominance of spiked structures with an angle less than 45^°^*C* which points toward the direction of cell motility (Figure 6g). This implies that the actin anchors contribute to enhanced motility in the grooves (Figure 6h).

**Figure 6.**
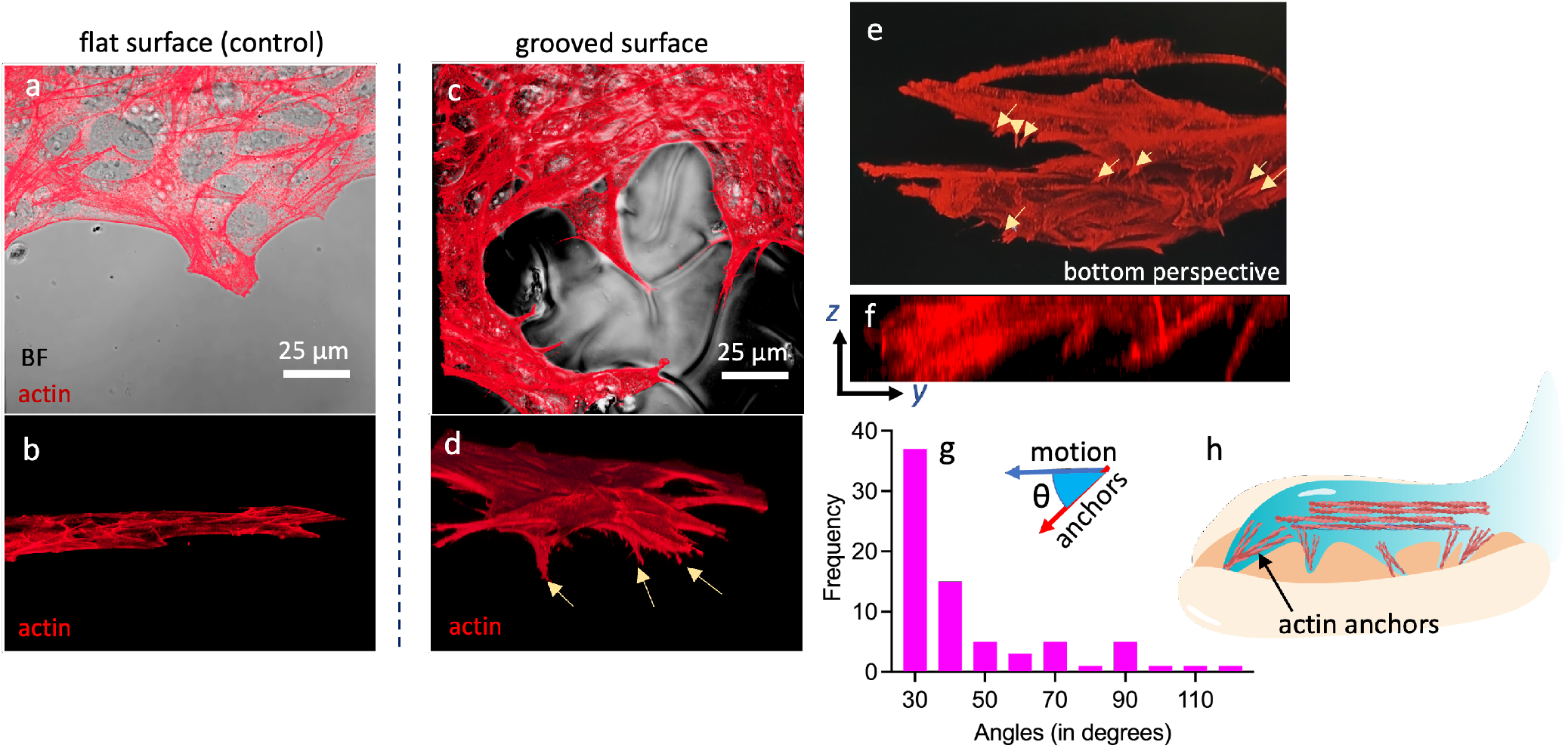
3D Actin protrusions in grooves and directionality of steering cell migration. (a) 2D merged image of BF and actin showing blunt leading edges in flat and soft hydrogel (control) surface. (b) 3D rendered side view perspective of cells on a flat hydrogel surface showing a very thin layer of cells with no prominent active bundling. (c) Merged BF and actin image of cells migrating on grooved hydrogel showing leading edges concentrating in the grooves. (d) 3D rendered side view perspective of cellular actin. The yellow arrows show typical anchor-like processes buried into the grooves of the hydrogel. (e) the bottom view perspective showing the anchor-like processes along the grooves which are aligned in different directions. (f) cross-section view of the leading edge of a cell inside a groove showing multiple anchors. (g) shows a histogram of anchor angles of actin embedded in the grooves with respect to the direction of motion. (h) in addition to thick actin bundles, cells in grooves develop multiple actin anchors into the groove, which promotes cell motility.

## DISCUSSION

Cell migration is an intelligent process involving a series of sensing and reacting activities in the body. When cells sense a topological attribute, they modulate their migratory process to adapt to their environment. In this article, we generated randomly oriented microgrooves that posed a unique challenge for the cells to navigate across. Since the hydrogel has ECM properties with RGD motifs recognizable by the cells, it closely mimics an ECM-like environment. As cells encounter grooved topologies, they sense the terrain, reorganize their cytoskeletal proteins, and attempt to partially explore potential paths before finalizing their preferred path. This touch-and-choose behavior is indicative of a form of cell migratory intelligence, though the factors influencing their choice/decision-making remain unknown.

In general, randomly oriented grooves have been shown to reduce cell motility, as they present a combination of groove directions. Grooves aligned with cell polarity increase cell speed by a factor of two, while grooves against cell polarity reduce it by a factor of eight. Micro-nano patterns have been reported to show similar behavior of cells migrating in the directions along and against cells’ polarity^77,78^. This suggests that the overall effect of such groove textures is a reduction in cell motility, even when the number of aligned and opposing grooves is balanced. Additionally, bifurcations in the grooves trigger the touch-and-choose mechanism, requiring cells to rearrange their cytoskeleton and select a path. A higher number of bifurcations of groove in the hydrogel results in further delayed cell motility. Although actin polymerization occurs at a fast pace of 500 nm/s, the depolymerization time is relatively slower (2 nm/s)^79^. These actin polymerization and depolymerization cycles are energy-intensive and consume significant amounts of Adenosine triphosphate (ATP) from the cell reservoir^80^. The cells at these bifurcations inadvertently need to spend substantial energy to proceed forward, further contributing to the delay. Thus, a groove texture can be seen as a tool that regulates cell motility based on the amount of ATP expenditure it causes.

To understand how the cells are propelling faster in the grooves aligned to cells’ polarity, our findings demonstrated that the effect is not solely due to the alignment of actin fibers in the direction of migration. Instead, it is a combinatorial approach. In addition to the thick bundling of actin fibers along the axis of cell polarity, cells in aligned grooves also form angular actin spikes that anchor and support their movement. These act as additional steering mechanisms, justifying the increased speed of cell motility in aligned hydrogel grooves. This closely resembles the skiing sports activity, where the skier uses ski poles to modulate speed and direction. The cells moving in these grooves are often forced to take sharp turns, curves, and bends. This anchoring activity may contribute to steering the cells in the directions within the grooves.

The groove textures act as guides, directing cells toward or away from specific paths. Depending on the orientation of the grooves relative to cell polarity, the speed of the cells can be modulated by a factor of hundred. This has several real-life applications. Materials can be innovatively designed with topological features to redirect cells to a desired track. For example, in wound healing applications, scaffolds or therapeutic patches can be designed to reintroduce or guide healthy cells from adjacent tissues to the wound site, promoting faster healing^81^. Hydrogels, known for their effectiveness in scarless wound healing, maintain moisture and can release therapeutic agents gradually. In addition, structural patterning on such hydrogels can provide important structural and topological aspects that enhance faster wound healing with reduced scarring. Conversely, scaffolds used in post-operative sites after tumor resections and biopsy can incorporate topological features that slow down the migration of potentially cancerous cells, reducing the risk of metastasis^82^. Further, topological features can be engineered to have multi-hierarchical complex topological features to trap cells in grooved traps that stop them from reentering the body^83–86^. For instance, a therapeutic patch can direct cancer cells toward a therapeutic compartment that can trap and kill these highly motile cells. This is particularly relevant in carcinoma^87,88^, where cancer cells gain motility through epithelial-to-mesenchymal transition^72,73,89^. By using chemoattractants, motile cancer cells could be guided via the grooves toward compartments equipped to trap and kill them.

## CONCLUSION

This study reveals the intricate relationship between cellular motility and substrate topology, emphasizing how cells navigate grooved environments by dynamically reorganizing their cytoskeleton, particularly actin. When grooves align with the cell’s polarity, migration speeds increase significantly, while misaligned grooves slow down movement and lead to the formation of additional podial structures. This demonstrates the critical role groove orientation plays in directing cellular behavior.

A key observation is the formation of actin-based anchoring structures within the grooves, which help steer and propel cells forward, especially in grooves aligned with their natural direction. This anchoring mechanism enhances traction and allows cells to navigate complex topologies, much like how a skier uses poles for control on challenging terrain. The study also identified a “touch-and-choose” behavior, where cells explore bifurcations before selecting a path—a process that, although energy-intensive, reflects a form of migratory intelligence.

These findings have broad potential applications, particularly in fields like wound healing and cancer treatment. Engineered grooved hydrogels could guide cellular behavior, accelerating cell migration for faster tissue repair or decelerating and trapping cancer cells to prevent metastasis. The ability to manipulate groove patterns offers promising possibilities for creating therapeutic scaffolds that control cell movement, opening new avenues for tissue engineering and regenerative medicine.

## MATERIALS AND METHODS

### Hydrogel preparation

GelMA was synthesized in-house as previously described^90,91^, exhibiting a high degree of substitution (97%). A solution of 50*mg/ml* LAP photoinitiator was prepared by dissolving it in sterile PBS pre-warmed to 50^°^*C*. To ensure homogeneous dissolution of LAP, the mixture underwent pulse ultrasonication for 5 to 10 minutes, followed by maintenance at 50^°^*C* in a water bath. The GelMA solution was fully hydrated in a water bath heated to 50^°^*C* for at least 2 hours. The GelMA precursor solution was prepared by adding the photoinitiator to occupy the remaining 10% volume of the GelMA solution, resulting in a final LAP concentration of 5*mg/ml* (or 0.5%*w/v*). The precursor was gently mixed by pipetting to prevent bubble formation. Subsequently, the precursor solution was transferred to a 37^°^*C* incubator to reduce viscosity and facilitate easy spreading before UV cross-linking.

### Preparation of spontaneously formed grooved hydrogel

The samples were made in 35 mm glass bottom Petri dishes with a center cover slip diameter of 20 mm. To create the texture, it is crucial that the surface is hydrophilic, such that the GelMA precursor forms an acute contact angle of about 30 degrees or less. This was achieved by either plasma coating the surface for 10 minutes. The next steps are done within the same day. Inducing surface hydrophilicity can be alternatively achieved by coating the glass surface with poly-L-lysine or by acid washing (1M HCL, 50 ^°^C, 4 hours). In these cases, it is important that the surface is completely dried (air dried at least for 2 hours) before progressing to subsequent stages. 25*mg/ml* of GelMA precursor (97% degree of substitution) was made with the final LAP concentration of 5*mg/ml* in PBS with pH 7.0. 100*µl* of the GelMA precursor solution was drop cast in the center of the glass bottom dish to create a well on the edges of the glass bottom well to seed cells later. UV cross-linking of GelMA was performed using a UV LED lamp (low power) with a central wavelength of 365nm, with exposure of 60 seconds. As the edges of the GelMA spot is thinner, the edges and the center of the hydrogel cures differently. The difference in the curing creates folding of the GelMA at the edges. Pre-warmed PBS at 37^°^*C* was used to immerse the photocured GelMA and kept in an incubator for 10 minutes. This generated spontaneous grooved structures at the edges of the hydrogel with a thickness of 200-300 *µ*m and a mean groove thickness of 2*µ*m. The size of the groove and the thickness of the groove textured patch can be modulated by changing the time of the photo-crosslinking (longer time led to making thicker patches of textured groove) and contact angle (lower contact angle makes thicker patches). The groove texture was observed while using GelMA concentration between 20 to 30 mg/ml. For creating textures on higher concentrations of GelMA a substantially longer curing time is needed (10 minutes for 50mg/ml GelMA). Subsequently, the PBS was aspirated to wash out any uncrosslinked GelMA and photo-initiator. The sample was kept hydrated in PBS in a cell culture incubator to prepare it for seeding fresh cells.

### Cell culture and seeding

Immortalized mouse embryonic fibroblasts were used between passages 3 and 10. The cells were maintained in DMEM high glucose media supplemented with 10% FBS and 1% antibiotic. During the experiment, the cells were maintained in a 37^°^*C* incubator with 5%*CO*_2_. Right before seeding the cells, the PBS was aspirated completely. 50,000 cells were seeded in the well next to the grooved hydrogel edges. We ensure that the drop does not touch the hydrogel edge at the beginning of the experiment. The dishes were allowed to incubate for 1 hour for cell attachment. After 1 hour, the unattached cells were washed out with PBS, and 2ml of fresh media was added to the dishes. After 24 hours when the cells were climbing the texture, the events were imaged with either label-free microscopy or fixed at desired time-points for further labeling and investigation.

### Label-free imaging

For studying the motility of the cells live^92^, we tracked the cells under a differential interference contrast (DIC) microscope for a period of up to 60 hours. The label-free DIC imaging was done in the GE Delta Vision microscope, using the objective lens with 10x, 0.40 Numerical Aperture (NA). To quantify the changes in the label-free intrinsic cellular properties, gradient light interference microscopy (GLIM) was performed. We used the Nikon Ti2E microscope along with the GLIM module by Phi Optics.

### Cell fixation and fluorescence labeling

The cells were fixed with 4% paraformaldehyde in PBS for 20 mins. For labelling the cells with fluorescent dyes, the cells were permeabilized in 0.3% Triton X-100 permeabilization buffer with 5% goat serum. Rabbit monoclonal to Vinculin (EPR8185, Abcam) was used as primary antibody, which was incubated with the sample overnight at 4^°^C. Goat anti-Rabbit Secondary Antibody was used tagged with Alexa Fluor 488 (A-11008, ThermoFisher). For labelling the F-actin, phalloidin-Atto 647 was used.

### Confocal microscopy

The confocal imaging for measuring the actin distribution was performed with Zeiss LSM800 confocal laser scanning microscope with a 40x 1.2 NA water immersion objective lens. The pinhole of 1 airy unit and a laser power of 0.5% was used. Correlative brightfield imaging was performed to identify actin structures in the grooved areas of the hydrogel.

### Image and data analysis

All the image analyses was done in Fiji software. Just Another Co-localization Plugin (JACoP) was used for measuring the co-localization between atin and vinculin. Groove thickness and groove angles was measured from phase contrast images. For generating standard deviation maps, Z-project function was used on the time lapse data of desired time windows. Temporal-Color Code function was used for temporally coding the data to determine active sites on the cell migration using time-lapse data in 20 hour windows. For SRRF super resolution microscopy processing, Nano-J SRRF plugin was used. GraphPad Prism 10 was used for statistical analysis.

## ACKNOWLEDGEMENTS

The research is funded by the following projects: Research Council of Norway projects-”Cyto-Motility and Cyto-Plasticity in Vitro Live-Cell Assay” (345442); Universitetet i Tromsø (UiT) grants-UiT Talent Supplementary and UiT Talent Innovation Main Project “CYMOPLIVE”; H2020 FET-Open RIA project “OrganVision” (964800) and ERC Starting Grant (804233). We also acknowledge UiT The Arctic University of Norway for funding article processing charges of the published article.

## AUTHOR CONTRIBUTIONS

B.G. conceptualized the study, wrote the paper, and performed the experiments. K.A. provided infrastructure for biological studies. All authors reviewed and revised the manuscript.

## References

1. Park, J. et al. Directed migration of cancer cells guided by the graded texture of the underlying matrix. Nat. materials 15, 792–801 (2016).

2. Heath, J. & Dunn, G. Cell to substratum contacts of chick fibroblasts and their relation to the microfilament system. a correlated interference-reflexion and high-voltage electron-microscope study. J. cell science 29, 197–212 (1978).

3. Olesen, S.-P., Claphamt, D. & Davies, P. Haemodynamic shear stress activates a k+ current in vascular endothelial cells. Nature 331, 168–170 (1988).

4. Hynes, R. O. The emergence of integrins: a personal and historical perspective. Matrix biology: journal Int. Soc. for Matrix Biol. 23, 333 (2004).

5. Kaverina, I., Krylyshkina, O. & Small, J. V. Microtubule targeting of substrate contacts promotes their relaxation and dissociation. The J. cell biology 146, 1033–1044 (1999).

6. Paul, C. D., Mistriotis, P. & Konstantopoulos, K. Cancer cell motility: lessons from migration in confined spaces. Nat. reviews cancer 17, 131–140 (2017).

7. Ghosh, B. et al. Arecanut-induced fibrosis display dual phases of reorganising glycans and amides in skin extracellular matrix. Int. J. Biol. Macromol. 185, 251–263 (2021).

8. Wolf, K., Muller, R., Borgmann, S., Brocker, E.-B. & Friedl, P. Amoeboid shape change and contact guidance: T-lymphocyte crawling through fibrillar collagen is independent of matrix remodeling by mmps and other proteases. Blood 102, 3262–3269 (2003).

9. Renkawitz, J. et al. Nuclear positioning facilitates amoeboid migration along the path of least resistance. Nature 568, 546–550 (2019).

10. Castro-Castro, A. et al. Cellular and molecular mechanisms of mt1-mmp-dependent cancer cell invasion. Annu. review cell developmental biology 32, 555–576 (2016).

11. Hanumantharao, S. N., Que, C. A., Vogl, B. J. & Rao, S. Engineered three-dimensional scaffolds modulating fate of breast cancer cells using stiffness and morphology related cell adhesion. IEEE Open J. Eng. Medicine Biol. 1, 41–48 (2020).

12. Agarwal, K. et al. Additive manufacturing enabled by electrospinning for tougher bio-inspired materials. Adv. Mater. Sci. Eng. 2018, 8460751 (2018).

13. Agarwal, K., Sahay, R. & Baji, A. Tensile properties of composite reinforced with three-dimensional printed fibers. Polymers 12, 1089 (2020).

14. Weigelin, B., Bakker, G.-J. & Friedl, P. Intravital third harmonic generation microscopy of collective melanoma cell invasion: Principles of interface guidance and microvesicle dynamics. IntraVital 1, 32–43 (2012).

15. Vasudevan, J., Lim, C. T. & Fernandez, J. G. Cell migration and breast cancer metastasis in biomimetic extracellular matrices with independently tunable stiffness. Adv. Funct. Mater. 30, 2005383 (2020).

16. Kumar, N. et al. Multilayered “smart” hydrogel systems for on-site drug delivery applications. J. Drug Deliv. Sci. Technol. 80, 104111 (2023).

17. Ghosh, B. & Agarwal, K. Gelma hydrogel mechanics affect the collective migration of fibroblast cells. In TISSUE ENGINEERING PART A, vol. 29 (MARY ANN LIEBERT, INC 140 HUGUENOT STREET, 3RD FL, NEW ROCHELLE, NY 10801 USA, 2023).

18. Cameron, A. R., Frith, J. E., Gomez, G. A., Yap, A. S. & Cooper-White, J. J. The effect of time-dependent deformation of viscoelastic hydrogels on myogenic induction and rac1 activity in mesenchymal stem cells. Biomaterials 35, 1857–1868 (2014).

19. Young, D. A., Choi, Y. S., Engler, A. J. & Christman, K. L. Stimulation of adipogenesis of adult adipose-derived stem cells using substrates that mimic the stiffness of adipose tissue. Biomaterials 34, 8581–8588 (2013).

20. Engler, A. J. et al. Embryonic cardiomyocytes beat best on a matrix with heart-like elasticity: scar-like rigidity inhibits beating. J. cell science 121, 3794–3802 (2008).

21. Hazeltine, L. B. et al. Temporal impact of substrate mechanics on differentiation of human embryonic stem cells to cardiomyocytes. Acta biomaterialia 10, 604–612 (2014).

22. Gershlak, J. R. et al. Mesenchymal stem cells ability to generate traction stress in response to substrate stiffness is modulated by the changing extracellular matrix composition of the heart during development. Biochem. biophysical research communications 439, 161–166 (2013).

23. Farouz, Y., Chen, Y., Terzic, A. & Menasché, P. Concise review: growing hearts in the right place: on the design of biomimetic materials for cardiac stem cell differentiation. Stem Cells 33, 1021–1035 (2015).

24. Engler, A. J. et al. Myotubes differentiate optimally on substrates with tissue-like stiffness: pathological implications for soft or stiff microenvironments. The J. cell biology 166, 877–887 (2004).

25. Hadden, W. J. et al. Stem cell migration and mechanotransduction on linear stiffness gradient hydrogels. Proc. Natl. Acad. Sci. 114, 5647–5652 (2017).

26. Ganguly, K. et al. Transcriptomic changes toward osteogenic differentiation of mesenchymal stem cells on 3d-printed gelma/cnc hydrogel under pulsatile pressure environment. Adv. Healthc. Mater. 12, 2202163 (2023).

27. Günay, K. A. et al. Myoblast mechanotransduction and myotube morphology is dependent on bag3 regulation of yap and taz. Biomaterials 277, 121097 (2021).

28. Engler, A. J., Sen, S., Sweeney, H. L. & Discher, D. E. Matrix elasticity directs stem cell lineage specification. Cell 126, 677–689 (2006).

29. Guvendiren, M. & Burdick, J. A. Stiffening hydrogels to probe short-and long-term cellular responses to dynamic mechanics. Nat. communications 3, 792 (2012).

30. Ploeg, M. C. et al. Culturing of cardiac fibroblasts in engineered heart matrix reduces myofibroblast differentiation but maintains their response to cyclic stretch and transforming growth factor β 1. Bioengineering 9, 551 (2022).

31. Gandin, A. et al. Simple yet effective methods to probe hydrogel stiffness for mechanobiology. Sci. reports 11, 22668 (2021).

32. Rajendran, A. K. et al. Trends in mechanobiology guided tissue engineering and tools to study cell-substrate interactions: a brief review. Biomater. Res. 27, 55 (2023).

33. Flynn, L. The use of decellularized adipose tissue to provide an inductive microenvironment for the adipogenic differentiation of human adipose-derived stem cells. Biomaterials 31, 4715–4724 (2010).

34. Xie, H., Cao, T., Franco-Obregón, A. & Rosa, V. Graphene-induced osteogenic differentiation is mediated by the integrin/fak axis. Int. J. Mol. Sci. 20, 574 (2019).

35. Harris, A. K., Wild, P. & Stopak, D. Silicone rubber substrata: a new wrinkle in the study of cell locomotion. Science 208, 177–179 (1980).

36. de Oliveira, N. B. et al. Natural membrane differentiates human adipose-derived mesenchymal stem cells to neurospheres by mechanotransduction related to yap and amot proteins. Membranes 11, 687 (2021).

37. Singhvi, R. et al. Engineering cell shape and function. Science 264, 696–698 (1994).

38. Kilian, K. A., Bugarija, B., Lahn, B. T. & Mrksich, M. Geometric cues for directing the differentiation of mesenchymal stem cells. Proc. Natl. Acad. Sci. 107, 4872–4877 (2010).

39. Wei, Q. et al. Bmp-2 signaling and mechanotransduction synergize to drive osteogenic differentiation via yap/taz. Adv. Sci. 7, 1902931 (2020).

40. Abagnale, G. et al. Surface topography enhances differentiation of mesenchymal stem cells towards osteogenic and adipogenic lineages. Biomaterials 61, 316–326 (2015).

41. Sonam, S., Sathe, S. R., Yim, E. K., Sheetz, M. P. & Lim, C. T. Cell contractility arising from topography and shear flow determines human mesenchymal stem cell fate. Sci. reports 6, 20415 (2016).

42. Fu, J. et al. Mechanical regulation of cell function with geometrically modulated elastomeric substrates. Nat. methods 7, 733–736 (2010).

43. Dalby, M. J. et al. Osteoprogenitor response to semi-ordered and random nanotopographies. Biomaterials 27, 2980–2987 (2006).

44. Zhou, K. et al. Nanoscaled and microscaled parallel topography promotes tenogenic differentiation of asc and neotendon formation in vitro. Int. journal nanomedicine 3867–3881 (2018).

45. Tay, C. Y. et al. Micropatterned matrix directs differentiation of human mesenchymal stem cells towards myocardial lineage. Exp. cell research 316, 1159–1168 (2010).

46. Morez, C. et al. Enhanced efficiency of genetic programming toward cardiomyocyte creation through topographical cues. Biomaterials 70, 94–104 (2015).

47. Takada, T. et al. Aligned human induced pluripotent stem cell-derived cardiac tissue improves contractile properties through promoting unidirectional and synchronous cardiomyocyte contraction. Biomaterials 281, 121351 (2022).

48. Hermans, L. et al. Scaffold geometry-imposed anisotropic mechanical loading guides the evolution of the mechanical state of engineered cardiovascular tissues in vitro. Front. bioengineering biotechnology 10, 796452 (2022).

49. Melo-Fonseca, F. et al. Reengineering bone-implant interfaces for improved mechanotransduction and clinical outcomes. Stem Cell Rev. Reports 16, 1121–1138 (2020).

50. Fiedler, J. et al. The effect of substrate surface nanotopography on the behavior of multipotnent mesenchymal stromal cells and osteoblasts. Biomaterials 34, 8851–8859 (2013).

51. McNamara, L. E. et al. Skeletal stem cell physiology on functionally distinct titania nanotopographies. Biomaterials 32, 7403–7410 (2011).

52. Sjöström, T. et al. Fabrication of pillar-like titania nanostructures on titanium and their interactions with human skeletal stem cells. Acta biomaterialia 5, 1433–1441 (2009).

53. von der Mark, K., Bauer, S., Park, J. & Schmuki, P. Another look at “stem cell fate dictated solely by altered nanotube dimension”. Proc. Natl. Acad. Sci. 106, E60–E60 (2009).

54. Lavenus, S. et al. Adhesion and osteogenic differentiation of human mesenchymal stem cells on titanium nanopores. Eur Cell Mater 22, 84–96 (2011).

55. de Peppo, G. M. et al. Osteogenic response of human mesenchymal stem cells to well-defined nanoscale topography in vitro. Int. journal nanomedicine 2499–2515 (2014).

56. Dalby, M. J. et al. The control of human mesenchymal cell differentiation using nanoscale symmetry and disorder. Nat. materials 6, 997–1003 (2007).

57. Wang, J. R. et al. Nanotopology potentiates growth hormone signalling and osteogenesis of mesenchymal stem cells. Growth Horm. & IGF Res. 24, 245–250 (2014).

58. Yang, J. et al. Nanotopographical induction of osteogenesis through adhesion, bone morphogenic protein cosignaling, and regulation of micrornas. ACS nano 8, 9941–9953 (2014).

59. Tsimbouri, P. M. et al. A genomics approach in determining nanotopographical effects on msc phenotype. Biomaterials 34, 2177–2184 (2013).

60. Poudineh, M. et al. Three-dimensional nanostructured architectures enable efficient neural differentiation of mesenchymal stem cells via mechanotransduction. Nano letters 18, 7188–7193 (2018).

61. Yim, E. K., Pang, S. W. & Leong, K. W. Synthetic nanostructures inducing differentiation of human mesenchymal stem cells into neuronal lineage. Exp. cell research 313, 1820–1829 (2007).

62. Yang, K. et al. Nanotopographical manipulation of focal adhesion formation for enhanced differentiation of human neural stem cells. ACS applied materials & interfaces 5, 10529–10540 (2013).

63. Ankam, S. et al. Substrate topography and size determine the fate of human embryonic stem cells to neuronal or glial lineage. Acta biomaterialia 9, 4535–4545 (2013).

64. Wang, B., Cai, Q., Zhang, S., Yang, X. & Deng, X. The effect of poly (l-lactic acid) nanofiber orientation on osteogenic responses of human osteoblast-like mg63 cells. J. mechanical behavior biomedical materials 4, 600–609 (2011).

65. Hu, J., Liu, X. & Ma, P. X. Induction of osteoblast differentiation phenotype on poly (l-lactic acid) nanofibrous matrix. Biomaterials 29, 3815–3821 (2008).

66. Liu, W. et al. Lower extent but similar rhythm of osteogenic behavior in hbmscs cultured on nanofibrous scaffolds versus induced with osteogenic supplement. Acs Nano 7, 6928–6938 (2013).

67. Cardwell, R. D., Dahlgren, L. A. & Goldstein, A. S. Electrospun fibre diameter, not alignment, affects mesenchymal stem cell differentiation into the tendon/ligament lineage. J. tissue engineering regenerative medicine 8, 937–945 (2014).

68. Kang, X. et al. Adipogenesis of murine embryonic stem cells in a three-dimensional culture system using electrospun polymer scaffolds. Biomaterials 28, 450–458 (2007).

69. El Khatib, M. et al. Electrospun plga fiber diameter and alignment of tendon biomimetic fleece potentiate tenogenic differentiation and immunomodulatory function of amniotic epithelial stem cells. Cells 9, 1207 (2020).

70. Chun, Y. W. et al. Combinatorial polymer matrices enhance in vitro maturation of human induced pluripotent stem cell-derived cardiomyocytes. Biomaterials 67, 52–64 (2015).

71. SenGupta, S., Parent, C. A. & Bear, J. E. The principles of directed cell migration. Nat. Rev. Mol. Cell Biol. 22, 529–547 (2021).

72. Mandal, M. et al. Modeling continuum of epithelial mesenchymal transition plasticity. Integr. Biol. 8, 167–176 (2016).

73. Mandal, M., Ghosh, B., Rajput, M. & Chatterjee, J. Impact of intercellular connectivity on epithelial mesenchymal transition plasticity. Biochimica et Biophys. Acta (BBA)-Molecular Cell Res. 1867, 118784 (2020).

74. Ghosh, B., Agarwal, K., Habib, A., Agarwal, K. & Melandso, F. Label-free acoustic and optical microscopy of live tumor spheroids in hydrogel for high-throughput 3d in-vitro drug screening. bioRxiv 2024–08 (2024).

75. Ghosh, B. et al. Molecular histopathology of matrix proteins through autofluorescence super-resolution microscopy. Sci. Reports 14, 10524 (2024).

76. Ghosh, B. & Agarwal, K. Viewing life without labels under optical microscopes. Commun. Biol. 6, 559 (2023).

77. Sun, L. et al. Controlling growth and osteogenic differentiation of osteoblasts on microgrooved polystyrene surfaces. PLoS One 11, e0161466 (2016).

78. Ramirez-San Juan, G., Oakes, P. & Gardel, M. Contact guidance requires spatial control of leading-edge protrusion. Mol. biology cell 28, 1043–1053 (2017).

79. Mogilner, A. & Oster, G. Force generation by actin polymerization ii: the elastic ratchet and tethered filaments. Biophys. journal 84, 1591–1605 (2003).

80. Atkinson, S. J., Hosford, M. A. & Molitoris, B. A. Mechanism of actin polymerization in cellular atp depletion. J. Biol. Chem. 279, 5194–5199 (2004).

81. Ghosh, B., Mandal, M., Mitra, P. & Chatterjee, J. Structural mechanics modeling reveals stress-adaptive features of cutaneous scars. Biomech. modeling mechanobiology 20, 371–377 (2021).

82. Bhowmik, A. et al. Portable, handheld, and affordable blood perfusion imager for screening of subsurface cancer in resource-limited settings. Proc. Natl. Acad. Sci. 119, e2026201119 (2022).

83. Agarwal, K., Sahay, R., Baji, A. & Budiman, A. S. Biomimetic tough helicoidally structured material through novel electrospinning based additive manufacturing. MRS Adv. 4, 2345–2354 (2019).

84. Budiman, A. S. et al. Impact-resistant and tough 3d helicoidally architected polymer composites enabling next-generation lightweight silicon photovoltaics module design and technology. Polymers 13, 3315 (2021).

85. Agarwal, K., Sahay, R., Budiman, A. & Baji, A. Development of strong and tough electrospun fiber-reinforced composites. Electrospun Polym. Compos. 287–313 (2021).

86. Budiman, A. S. et al. Modeling impact mechanics of 3d helicoidally architected polymer composites enabled by additive manufacturing for lightweight silicon photovoltaics technology. Polymers 14, 1228 (2022).

87. Ghosh, B. et al. Quantitative in situ imaging and grading of oral precancer with attenuation corrected-optical coherence tomography. Oral oncology 117, 105216 (2021).

88. Ghosh, B., Mandal, M., Mitra, P. & Chatterjee, J. Attenuation corrected-optical coherence tomography for quantitative assessment of skin wound healing and scar morphology. J. Biophotonics 14, e202000357 (2021).

89. Sarkar, A. et al. Autofluorescence signatures for classifying lung cells during epithelial mesenchymal transition. RSC advances 6, 77953–77962 (2016).

90. Ghosh, B., Fenton, K. A. & Agarwal, K. Mechano-chemical insights in diabetic kidney disease through 3d pathotypic model of mesangium. bioRxiv 2023–12 (2023).

91. Ghosh, B. & Agarwal, K. Kidney mesangium organotypic culture reveals co-stimulatory mechano-chemical interplay for fibrotic manifestation. Biophys. J. 123, 405a (2024).

92. Ghosh, B. & Chatterjee, J. Advances in medical imaging for wound repair and regenerative medicine. In Regenerative Medicine: Emerging Techniques to Translation Approaches, 57–76 (Springer, 2023).

